# vScreenML v2.0: Improved Machine Learning Classification for Reducing False Positives in Structure-Based Virtual Screening

**DOI:** 10.1101/2024.10.08.617248

**Authors:** Grigorii V. Andrianov, Emeline Haroldsen, John Karanicolas

## Abstract

Enthusiastic adoption of make-on-demand chemical libraries for virtual screening has highlighted the need for methods that deliver improved hit-finding discovery rates. Traditional virtual screening methods are often inaccurate, with most compounds nominated in a virtual screen not engaging the intended target protein to any detectable extent. Emerging machine learning approaches have made significant progress in this regard, including our previously-described tool vScreenML. Broad adoption of vScreenML was hindered by its challenging usability and dependencies on certain obsolete or proprietary software packages. Here, we introduce vScreenML 2.0 (https://github.com/gandrianov/vScreenML2) to address each of these limitations with a streamlined Python implementation. Through careful benchmarks, we show that vScreenML 2.0 outperforms other widely-used tools for virtual screening hit discovery.

## Introduction

Starting points for drug discovery often arise from screening small-molecule libraries. Hits from such collections are typically recognized either by explicitly evaluating each compound in a relevant functional assay (e.g., biochemical screening), by pulling out binders from a pool of barcoded compounds (e.g., DNA-encoded libraries), or by computationally selecting compounds that fit with the desired site on the target protein surface. Although traditional biochemical screening affords the most directly relevant readout, its application is typically limited to collections of about 2 million compounds. By contrast, pooled DNA-encoded libraries can comprise 1 billion compounds: however, many apparent hits do not show activity when re-synthesized without the fused DNA (for reasons that include mis-identification of the screening hit, binding that relies on the linked DNA, and “silent” binders that do not confer any effect on protein function).

In recent years, chemical vendors significantly increased size of their catalogs of compounds by enumerating specific building blocks that can be combined using robust chemical transformations. This approach can lead to enormous libraries of new compounds that are readily synthetically tractable: for example, Enamine compiled offerings of ∼29 billion “make-on-demand” compounds for purchase [1]. Even the most ambitious HTS campaigns cannot explicitly access these enormous “virtual” compound collections, however, which has amplified the importance of using computational screening methods.

Several studies have employed molecular docking to screen multi-million compound libraries, in some cases yielding advanceable hits. The fraction of computational hits that show activity is often dependent on the target class, with GPCR ligands often yielding high hit rates (and initial hits that can also be quite potent). Among GPCR targets, for instance, screening a virtual library of 75 million tetrahydropyridines against the serotonin 5-HT2A receptor prioritized 17 compounds for experimental validation: 4 of these (24%) proved to be active, with binding affinities in the low-micromolar range (0.67–3.9 μM) [2]. A study targeting the MT_1_ and MT_2_ receptors screened 150 million compounds, and found that 15 of 38 compounds tested (39%) were active in the nano- to micromolar range (from 1 nM to 15 μM) [3]. In another study, screening 138 million compounds against the D4 dopamine receptor found that 122 hits from among 549 tested (22%) with more than 50% inhibition at 10 μM [4]. A screen of 490 million compounds against the σ2 receptor found that 124 of 484 compounds tested (26%) were active in the nano- to micromolar range [5]. Screening over 300 million diverse molecules targeting the α2A-adrenergic receptor led to selection of 48 compounds, of which 30 compounds (63%) showed activities ranging from 1.7 nM to 9.4 μM [6]. Screening 115 million compounds against the CysLT receptor subtypes yielded 10 of 71 (14%) for CysLT1R and 25 of 68 (37%) for CysLT2R [7]. A focused screen of 140 million triazoles and isoxazoles against CB_2_ provided 11 hits, of which 6 compounds (55%) showed activity below 10 μM [8].

For non-GPCR targets, though, hit rates are typically lower. For example, screening 99 million compounds against AmpC β-lactamase yielded five active compounds (1.3–400 μM) from 44 tested (11%) [4]. A large-scale screen of 235 million compounds against the SARS-CoV-2 main protease (Mpro) provided 3 hits from 100 compounds tested in an enzyme activity assay (3%), with affinities between 23 and 61 μM [9]. A screen of 400 million lead-like molecules against Mac1 produced 13 hits from 124 selected compounds (10%), with IC_50_ values ranging from 42 to 504 μM [10]. Finally, screening 1.3 billion compounds against KEAP1 identified 69 hits with submicromolar binding affinity from 590 tested compounds (12%) [11].

Hit-finding discovery rates were also notably low in the Critical Assessment of Computational Hit-finding Experiments (CACHE) challenges [12]. In the first of these benchmarking initiatives, for example, 23 participants were offered the opportunity to each choose up to 100 compounds for testing against a target selected by the organizers (the WDR domain of LRRK2). When tested in a primary SPR assay, only 73 compounds (3.2%) showed any hint of activity, and many of these hits did not replicate in secondary assays. Subsequent (ongoing) benchmarks have also showed similar outcomes. It is worth noting that the organizers intentionally select challenging/unprecedented targets for these benchmarks, which likely contribute to the low hit rates.

As highlighted above then, most compounds selected in ultra-large virtual screening campaigns turn out to be false positives. While vendors such as Enamine have dramatically reduced the expense associated with procuring compounds prioritized by the computational screen, the cost of testing these candidates is still significant and scales with the number of compounds to be tested. Whereas false negatives in screening simply represent missed opportunities for alternate hits, false positives represent a very real expense because they consume wet-lab time and reagents. Even in the highly successful screens cited above, the vast majority of virtual screening hits are not active when characterized in biochemical assays (i.e., hit rates that are typically far below 50%).

To address this, we recently developed a machine learning classifier dubbed “vScreenML” [13]. This model was trained to distinguish structures of active complexes from carefully curated decoys that would otherwise represent likely false positives. After validating the model with a series of retrospective benchmarks, we applied vScreenML to select hits from an Enamine library docked against human acetylcholinesterase (AChE). The top 23 compounds were purchased and characterized in a biochemical assay: this experiment revealed that most of the compounds prioritized by vScreenML were indeed AChE inhibitors, with more than half showing IC_50_ lower than 50 μM and the best hit yielding a K_I_ value of 175 nM. Importantly, none of these hits bore any resemblance to known AChE inhibitors (and AChE had not been used in training), confirming that the model had not simply memorized interactions from training.

Despite the dramatic performance of vScreenML relative to standard approaches, certain hurdles slowed its widespread adoption. Specifically, it required complicated manual compilation of certain programs needed calculating features that describe each docked model, including multiple outdated or expensive dependencies that proved to be prohibitive for many users. Here, we therefore report an updated version called vScreenML 2.0 (https://github.com/gandrianov/vScreenML2). This update is far easier to install and use, and also avoids the dependencies that previously proved cumbersome. At the same time, we also updated the model by including newly-released structures from PDB and incorporating several additional features for enhanced discriminative power.

## Computational Approach

As with the original vScreenML approach [13], our new model is intended to distinguish active compounds from inactive compounds. In a virtual screening workflow, vScreenML 2.0 is thus intended to be used after the contents of a library have been docked to the target protein, for selecting candidate compounds that should be advanced for explicit testing in biochemical or cellular assays – without bringing forward an abundance of false positives (**Figure 1**).

**Figure 1:**
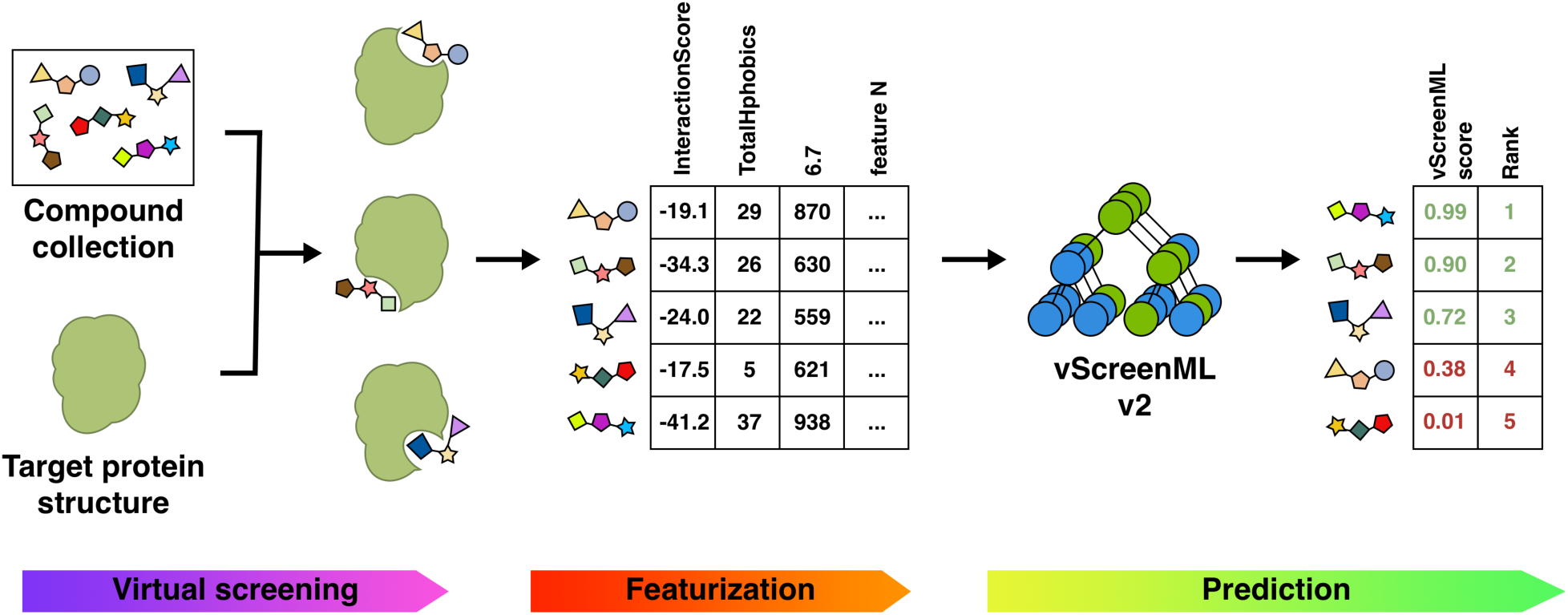
Incorporating vScreenML 2.0 into a virtual screening pipeline. Candidate protein-ligand complexes can be generated either by receptor-based methods (i.e., docking) or receptor-based methods (i.e., pharmacophoric alignment to one or more known ligands). Scoring each of the resulting complexes with vScreenML 2.0 involves extracting “features” from each model using the PyRosetta, Binana, RF-Score, RDKit, LUNA and PocketDruggability packages. These features are used as input to vScreenML 2.0, which uses an XGBoost-based model to produce a single score reflecting the prioritization of the compounds in the screening collection.

As described below, this updated version of vScreenML is more user-friendly for broad audiences, avoids using software with potentially costly licenses, and also adds flexibility that facilitates incorporation into complex virtual screening pipelines. To achieve these objectives, we replaced the C++-based Rosetta bundle with the Python-based PyRosetta package [14], we replaced MGLTools [15] with ODDT [16], we replaced ChemAxon cxcalc with the open-source RDKit equivalent [17], and we removed the dependency on OpenEye’s SZYBKI [18]. When training the new model, we also took this opportunity to incorporate newly-released structures in the PDB, and apply updated rules for defining which structures to include. Finally, we also incorporated new features to better capture several structural measures that were not well-described in the original model: buried unsatisfied polar groups, 2D structure properties of the ligand, and shape-based information from the protein pocket.

### Preparation of dataset (actives and decoys)

The original vScreenML approach [13] started from a set of (active) protein-ligand complexes from the RCSB Protein Data Bank (PDB) [19,20]. Each active compound was used to generate three property-matched “decoys” by selecting compounds with high 3D similarity to the corresponding active, from a set of physicochemically-matched candidates provided by the “Directory of Useful Decoys, Enhanced” (DUD-E) server [21]. The activity of these decoys have not been explicitly been tested for the corresponding protein target, but they are presumed to be inactive. Actives and decoys were refined using the same protocol, to avoid any clues from how the structures were prepared that could lead to artifactual performance by the classification model. The overall approach for training vScreenML 2.0 was conceptually similar, as summarized below (**Figure 2**).

**Figure 2:**
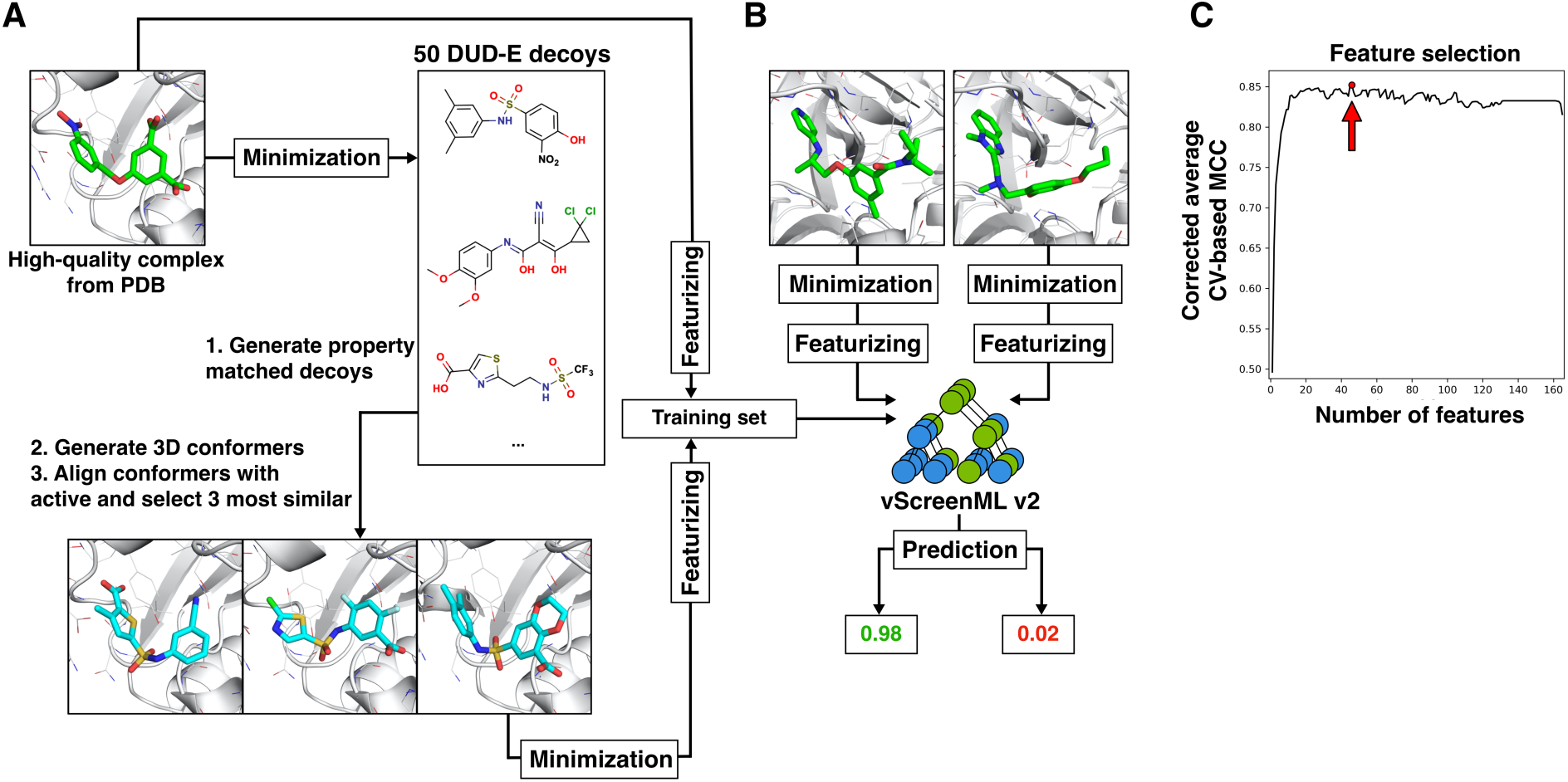
Dataset and training for vScreenML 2.0. **(A)** The dataset for training and testing comprises a set of active complexes, along with compelling decoy complexes built to match the active complexes. Actives were drawn from the Protein Data Bank (PDB), refined using Protoss, and then minimized using PyRosetta. Three decoy complexes were built from each active complex, by using ROCS to identify conformations of compounds from DUD-E that can adopt similar 3D structures as the active compound. Decoy complexes were refined the same way as the active complexes. **(B)** The structure of each active and decoy complex were used to calculate 166 numerical features: these were used to train the vScreenML 2.0 classifier. Actives were assigned label 1 and decoys were assigned label 0, so vScreenML 2.0 output scores range between 0 and 1. **(C)** To enhance robustness and avoid potential overtraining, feature reduction was applied. The final vScreenML 2.0 model includes only 49 of the original 166 features, since this model maximizes the Matthews correlation coefficient (MCC).

For vScreenML 2.0, we began with the updated contents of the PDB. We compiled a list of structures from the RCSB Protein Data Bank (PDB) that met the criteria: A) experimental method “X-Ray diffraction”, B) refinement resolution “< 2.5 Å”, C) and polymer entity type “Protein”. There were 132,146 PDB entries meeting these criteria. These structures were then filtered by physicochemical and structural features of ligand and protein-ligand complex presented in **Table S1**: specifically, these filters are primarily intended to restrict the training set actives to “drug-like” ligands (the intended domain of application for vScreenML 2.0) by removing the numerous non-drug-like biological ligands in the PDB (e.g. ATP, sugars, lipids, etc). Structures were also removed if the model quality was low, if it was incompletely resolved, or if it did not fit with the electron density.

The filtered data set comprised 1,407 PDB entries. For this collection, protonation states were assigned (along with placement of hydrogen atoms) using Protoss; however, 86 structures could not be refined, and were therefore excluded: this left a total of 1,321 unique PDB entries. Among these, some had multiple copies of the complex in the asymmetric unit. These copies have slight differences in the coordinates relative to one another, and thus differences in the calculated features as well. As a form of data augmentation, we retained these multiple copies of the same complex: we expected that this would help make vScreenML 2.0 resistant to noise in the training set. Thus, our set of actives comprised 1,806 instances, of which 1,321 are unique protein-ligand complexes. Each structure was minimized using PyRosetta prior to calculating vScreenML 2.0 features.

From each of the 1,806 actives, 3 property-matched decoys were also generated. To do so, the DUD-E server was first applied to each ligand in the collection of actives, leading to 50 candidate decoy ligands. These were then filtered based on 2D structural considerations used for the active ligands (**Table S1**), to ensure that there would be no systematic differences between active versus decoy ligands that could lead to spurious model performance. For each of the candidate decoys that passed the filters, 500 low-energy conformers were generated using OpenEye OMEGA [22] and these were aligned to the corresponding active structure using ROCS [23]. The three best-matched decoys (evaluated using TanimotoCombo score) were selected, because these would be most likely to fit with the protein pocket in a compelling manner. For certain actives, there were not three candidate decoys from DUD-E that passed these filters; in those cases, fewer than three decoys were generated. The alignment from ROCS was used to place the decoy ligands into the protein structure from the active ligand, then the model was refined using Protoss [24,25] and minimized with PyRosetta. Ultimately this procedure led to a total of 4,475 decoys, reflecting the fact that some of the 1,806 actives produced fewer than three decoys.

### Model training and feature selection

As with original vScreenML approach [13], numeric descriptors (features) were calculated from the structure of each protein-ligand complex. These were used in training the XGBoost [26] machine learning (ML) model, and when applying the model in practice. As noted earlier, some elements of original vScreenML features were not particularly user-friendly: these dependencies were eliminated in vScreenML 2.0, and a completely new Python-based framework was developed for vScreenML 2.0. Additionally, several new features are also included in this new implementation. A total of 166 features are calculated for each protein-ligand complex: these are reported in **Table S2**.

To prevent potential information leakage during training and testing, the dataset was divided into distinct groups by clustering protein sequences with MMSeq2 [27]. By ensuring that related proteins are always placed together in either the training set or the test set, we avoid potential overtraining in which the model sees a very similar complex in both training and testing. It is important to avoid this scenario, because it can lead to artificially inflated performance of the model and disappointing results in future prospective applications [28].

Clustering on protein sequences yielded 269 clusters that were used for cross-validation, by splitting the data into multiple sets for training and testing across several iterations. In each iteration, proteins from all clusters except one were used to train the model, while the remaining cluster was used for testing. This process was repeated until each cluster had been used as a test set. The final performance score of the model was obtained by averaging the performance metrics from all iterations, providing a more robust and reliable assessment of the model’s ability to generalize to unseen data.

To promote model generalization and avoid potential overfitting associated with using 166 features, we sought to eliminate features not contributing to model performance. We applied sequential backward floating feature selection, as implemented in the mlxtend [29] package in combination with cross-validation on the 269 protein sequence clusters. We found that the model which maximized the Matthews correlation coefficient required 49 features (**Figure 2c**): the features included in the final model are indicated in **Table S2**.

As with the original vScreenML implementation, we used the binary Extreme Gradient Boosting (XGB) framework [26] for our model. In model training, hyper-parameters of the XGB model (“n_estimators”, “max_depth” and “learning_rate”) were optimized using Optuna [30] to find the set of parameters that gave the best AUC upon 10-fold cross-validation.

### Benchmarking with DEKOIS2

Performance of vScreenML 2.0 was evaluated using the DEKOIS2 [31,32], a collection of active and decoy compounds for 81 protein targets. Models for the actives and the decoys in complex with their cognate target proteins were obtained from docked set created to evaluate the docking tool KarmaDock [33,34]. The structures were minimized using PyRosetta to make them more energetically consistent with the vScreenML 2.0 features.

To prevent any potential information leakage from training set, we clustered the protein targets from DEKOIS2 along with our training set, and excluded from the training set any targets with high sequence similarity to the DEKOIS2 protein targets.

The performance of vScreenML 2.0 was compared with other virtual screening tools that allow scoring in place of pre-generated poses, specifically AA-Score [35] and GNINA [36]. As a performance metric, we used enrichment factor of actives in top 1% (EF1%) of predicted scores. Statistical comparison of methods was carried out by removing “near ties” (absolute difference in EF1% less than 3), then using the binomial test implemented in the SciPy package [37].

### Data Availability

Minimized 3D structures of proteins in complex with actives and decoys, along with calculated features, are available in Zenodo (https://doi.org/10.5281/zenodo.10819385). The vScreenML 2.0 package and source code are located in GitHub (https://github.com/gandrianov/vScreenML2).

## Results

As noted above, vScreenML 2.0 reduces inconvenient software dependencies and also includes new features not present in the original implementation. These include ligand potential energy, buried unsatisfied atoms for select polar groups in ligand, additional 2D structural features of ligand, complete characterization of interface interactions in protein-ligand complex and pocket shape features (**Table S2**). To ensure model generalization (and avoid potential overtraining), we identified the 49 most important features to include in the model, rather than allow all 165 features (**Figure 2c**).

As a first evaluation of vScreenML 2.0, we compared its performance to that of the original version of vScreenML [13]. A direct comparison is complicated by the fact that the models were trained on different datasets. Each model should only be characterized using a held-out test set, and there is not a defined set common to either model that was not used in training. Nonetheless, when both models are applied to data from their respective datasets that was not used in training, the performance of vScreenML 2.0 far exceeds that of the original version (**Figure 3**).

**Figure 3:**
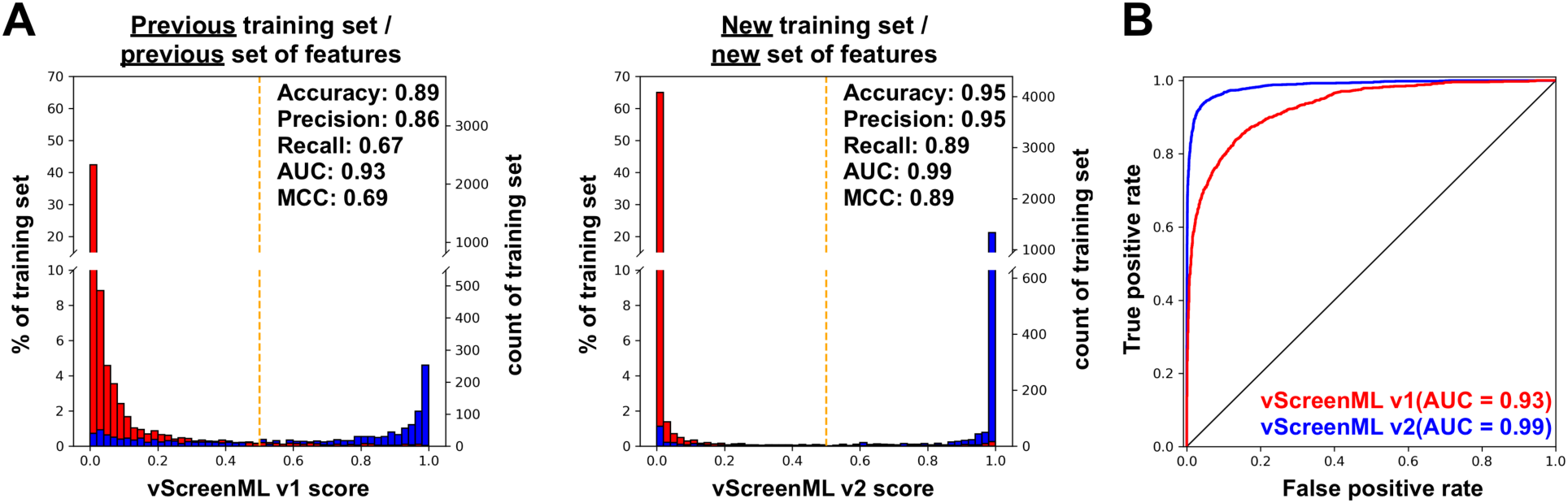
Performance of vScreenML on a test dataset. **(A)** Discriminating performance of the original vScreenML model (*left*) versus the new vScreenML 2.0 (*right*). The vScreenML score for each data point was calculated using cross-validation where each fold was prepared by clustering protein sequences using MMSeq2. **(B)** Receiver-operator characteristic plots showing performance of vScreenML 2.0 relative to the original model.

In each case, each of the models was applied to score protein-ligand complexes not used in training, involving exclusively protein targets dissimilar to those in the training set. Both versions of vScreenML recognized the active complexes with high scores (close to 1), and the decoy complexes with low scores (close to 0) (**Figure 3a**). The primary distinction that is evident between the two distributions is that vScreenML 2.0 mis-categorized fewer of the active complexes with low scores, compared to the original: this is reflected numerically in the higher recall value for vScreenML 2.0 (from 0.67 in the original to 0.89 in the new model). The improved recall, coupled with improved precision, leads to a dramatic improvement in Matthews correlation coefficient (MCC) for the new model (from 0.69 in the original to 0.89 in vScreenML 2.0). The improved performance is also dramatically evident when the same results are plotted as a receiver operating characteristic (ROC) curve (**Figure 3b**), demonstrating the enhanced performance of vScreenML 2.0 when applied to classification of held-out protein-ligand complexes.

Next, we compared the performance of vScreenML 2.0 in comparison with other widely-used tools for virtual screening hit discovery. As a representative empirical scoring function we selected AA-Score, a model that combines a broad variety of energetic components into a single score and showed impressive performance relative to a broad slate of other common scoring functions [35]. As a classic scoring function we used the GNINA [36] package’s implementation of AutoDock Vina [38], historically one of the most widely-used tools for virtual screening. Finally, as a representative of the deep learning models that have recently been described in the literature, we selected the GNINA’s CNNaffinity score [39]. This scoring method was shown to outperform AutoDock Vina scoring in a standard benchmark [40], though the authors acknowledge that some lingering bias (favoring GNINA’s CNNaffinity score) may come from recognizing property distributions rather than the details of molecular interactions.

We evaluated each of these methods using the DEKOIS2 dataset [31,32]. This benchmark comprises 81 protein targets, with the 2D chemical structures of 30-40 actives and 800-1200 property-matched decoys for each target. Each of these ligands were docked to the corresponding protein target using Schrödinger Glide in the context of a separate study [33,34], and so we re-used these starting poses. Each starting pose was minimized using PyRosetta to make them energetically consistent with the vScreenML 2.0 features.

While additional deep learning-based docking tools have also been developed for virtual screening hit discovery, such as KarmaDock [34] and RTMScore [41], they have seen most of the DEKOIS2 targets in the course of training: this makes it impossible to determine the extent to which their performance on this test set reflects the true performance that should be expected in a prospective application. This point is reinforced through the behavior of KarmaDock on PYGL ligands (**Table S3**). Structures of this target in the “in” conformation but not the “out” conformation were included in the KarmaDock training set, and we find that KarmaDock yields far superior performance for DEKOIS2 ligands in the “in” conformation. While GNINA’s CNNaffinity score was indeed trained on some of the DEKOIS2 targets, it saw fewer of these in training than KarmaDock or RTMScore (**Table S3**).

To avoid *any* potential for vScreenML 2.0 recognizing features of the DEKOIS2 targets, meanwhile, we used a model for each target that excluded the corresponding protein cluster from training (i.e., all data from any sequence-related proteins was left out of training).

To evaluate performance of the four selected scoring methods (vScreenML 2.0, AA-Score, AutoDock Vina, and GNINA CNNaffinity), we used each method to rank-order each of the pre-built DEKOIS2 models for each target, and calculated the enrichment factor of actives in top 1%. For a typical target with 30 actives and 970 decoys (1000 models), one would expect to find by random chance 0.3 actives in the top-scoring 10 models; a scoring tool that finds 3 actives within the top-scoring 10 models would therefore have an enrichment factor of 10.

The distribution of EF1% values for the 81 protein targets are first compared for vScreenML 2.0 and for AA-Score (**Figure 4a**, *top*). It is evident from the superposed histograms that AA-Score had more targets with very low EF1% values (0 to 7), whereas vScreenML 2.0 had more targets with higher EF1% values (10 and above). Extending the analysis to comparison of individual targets (**Figure 4a**, *bottom*), there are a few individual targets for which both methods perform well. For the most part, however, the clearest observation is that there are many more points above the diagonal than below, representing examples in which vScreenML 2.0 provides superior performance to AA-Score. Analysis using the binomial test confirms the statistical significance of the observation that vScreenML 2.0 is better for more targets than AA-Score (p < 9 x 10^-6^). The performance of AutoDock Vina was similarly surpassed by vScreenML 2.0 (p < 2 x 10^-9^) (**Figure 4b**).

**Figure 4:**
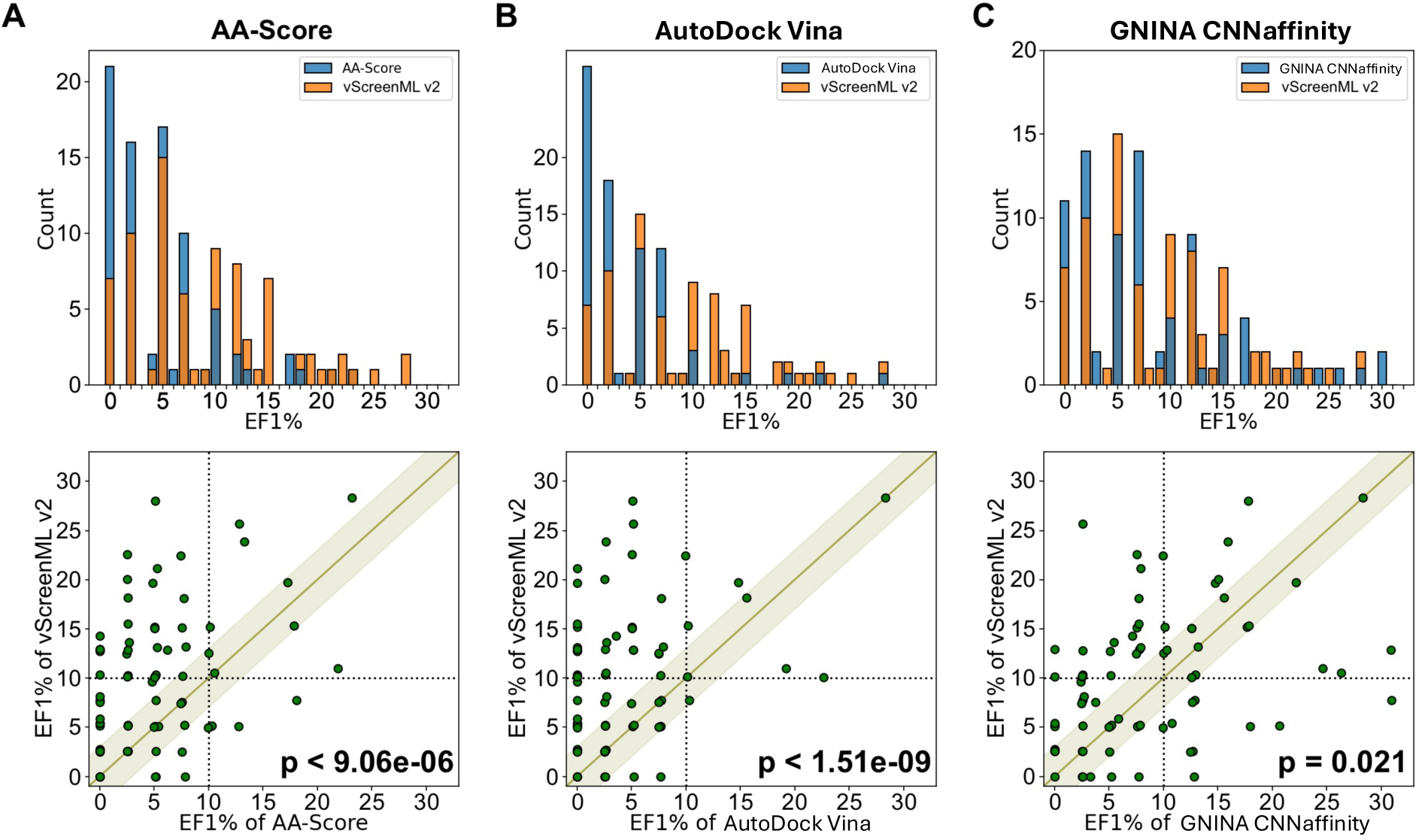
Comparison of vScreenML 2.0 to other modern scoring methods, using the pre-docked models from the DEKOIS2 benchmark. The test set consists of 81 target proteins, each of which is associated with 30-40 active compounds and 800-1200 of decoy compounds. The performance of vScreenML 2.0 is compared with that of **(A)** AA-Score, **(B)** AutoDock Vina, and **(C)** GNINA CNNaffinity. In each case, performance is characterized via the enrichment factor of actives in the top 1% (EF1%). *Top:* distribution of EF1% values across the set of 81 targets. *Bottom:* comparison of performance for specific targets. In all three cases, there are more points above the diagonal band than below the diagonal band, indicating that vScreenML 2.0 has superior performance on more targets. *P*-values was calculated using the binomial test (excluding points within the green band, for which the absolute difference in EF1% was less than 3).

Comparing vScreenML 2.0 to GNINA CNNaffinity showed more similar performance in this experiment. Superposition of the EF1% distributions shows slightly more low values for GNINA CNNaffinity, and slightly more high values for vScreenML 2.0 (**Figure 4c**, *top*). Comparing the 81 individual targets, both methods yielded similar performance for 35 targets (absolute difference in EF1% less than 3). Of the other 46 targets, vScreenML 2.0 was superior for 28 targets and GNINA CNNaffinity was superior for the other 18; per the binomial test, this difference is indeed statistically significant (p = 0.021) (**Figure 4c**, *bottom*). We also cannot fully rule out the possibility that GNINA CNNaffinity’s performance was bolstered by “recognizing” some interactions from its own training, which includes proteins from many of the target classes included in the DEKOIS2 benchmark. Nonetheless, there appears to be little agreement between which targets are “easiest” for these two methods: this may imply that the methods are recognizing different features in the active complexes, and that a consensus scoring approach may prove superior to either method alone.

## Discussion

vScreenML 2.0 addresses a key challenge in virtual screening, the high rate of false positives generated by traditional methods. The initial version of vScreenML proved extremely successful in prospective applications, but had strong drawbracks with respect to usability. These included a complex installation process, a complicated pipeline requiring manual intervention, and inclusion of multiple outdated, complex, or paid dependencies.

Through this update, we have addressed each of these issues by rewriting the entire framework in a more accessible programming language (Python); this both promotes ease of use and installation, while also enabling straightforward integration of vScreenML 2.0 into existing virtual screening pipelines. This changes thus opens the door for a broader user base, including non-experts who may have been previously deterred by the technical complexities of the earlier version. By improving the calculation pipeline and removing dependencies on obsolete or proprietary software, we have also made the tool more user-friendly and compatible with modern computational environments.

Finally, we updated the training set for the machine learning model by incorporating the latest crystal structures from the PDB, applying stringent new selection criteria to improve data quality, and introducing several new features that capture more nuanced aspects of protein-ligand interactions. Though we could not directly compare vScreenML 2.0 against the original version using the same test set, performance of the newer version appears superior. Nonetheless, the new model still remains to be validated in future prospective experiments to evaluate how these enhancements translate to meaningful gains in real-world applications.

Relative to standard empirical scoring functions such as AA-Score and AutoDock Vina, vScreenML 2.0 shows clear superiority for virtual screening hit discovery. While deep learning methods such as KarmaDock, RTMScore, and GNINA use CNN-based scoring functions that seem to provide similar or superior accuracy to vScreenML 2.0 in some respects, their ability to generalize to new targets remains unclear. Rigorous benchmarking can be difficult because these methods often incorporate all available public data in training, such that there is no single held-out test set that can be used to compare different methods. A key benefit of vScreenML is that the simplicity of the model and rigorous splitting between the training and test sets ensures generalizability to new protein target classes. As noted in the context of the DEKOIS2 benchmark, the vScreenML 2.0 performance presented above included no related proteins for each test set target, confirming generalization of the underlying models.

Looking ahead, we note that the size of ultra-large chemical libraries has grown much faster than the ability to explicitly dock all compounds in a modern library. To address this issue, new strategies have been employed to reduce the computational burden. One such approach entails using active learning together with docking results [42,43]. After explicitly docking a subset of the library against the target, ML models can be trained to rapidly identify compounds likely to yield good docking scores, without explicitly docking them. Using this model, a small subset of compounds can be prioritized for explicit docking from the vast chemical space available in the library. These top-ranked predictions are validated through explicit docking, the model can be retrained using this additional data and the process repeated.

We anticipate that vScreenML 2.0 may provide further value in these active learning frameworks. Specifically, the improved identification of likely binders may allow enhanced steering the search for compounds worth explicitly docking, in addition to guiding the final selection of compounds for experimental characterization.

## Acknowledgements

This work was supported by the W.M. Keck Foundation and by the NIH National Institute of General Medical Sciences (R01GM141513). This research was also funded in part through the NIH/NCI Cancer Center Support Grant P30CA006927.

This work used the Extreme Science and Engineering Discovery Environment (XSEDE) allocation MCB130049, which is supported by National Science Foundation grant number 1548562. This work also used computational resources through allocation MCB130049 from the Advanced Cyberinfrastructure Coordination Ecosystem: Services & Support (ACCESS) program, which is supported by National Science Foundation grants 2138259, 2138286, 2138307, 2137603, and 2138296.

We thank ChemAxon for providing an academic research license.

## Supporting Information

**Table S1.**
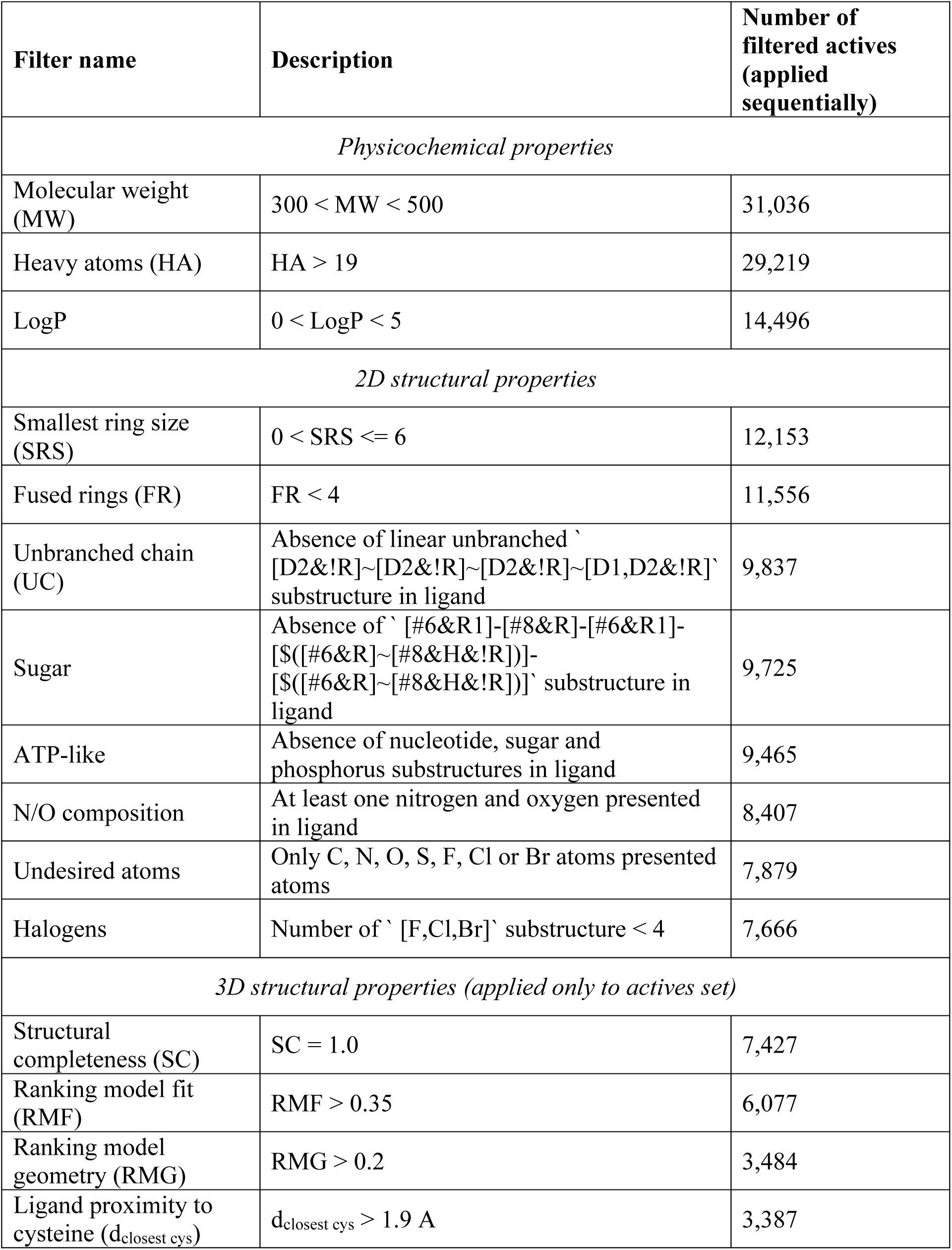
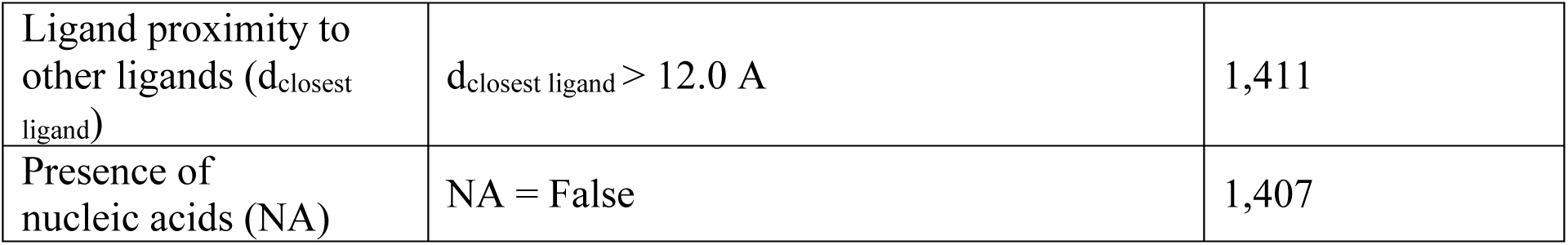
Filtering rules applied to molecules obtained from RCSB and DUD-E.

**Table S2.**
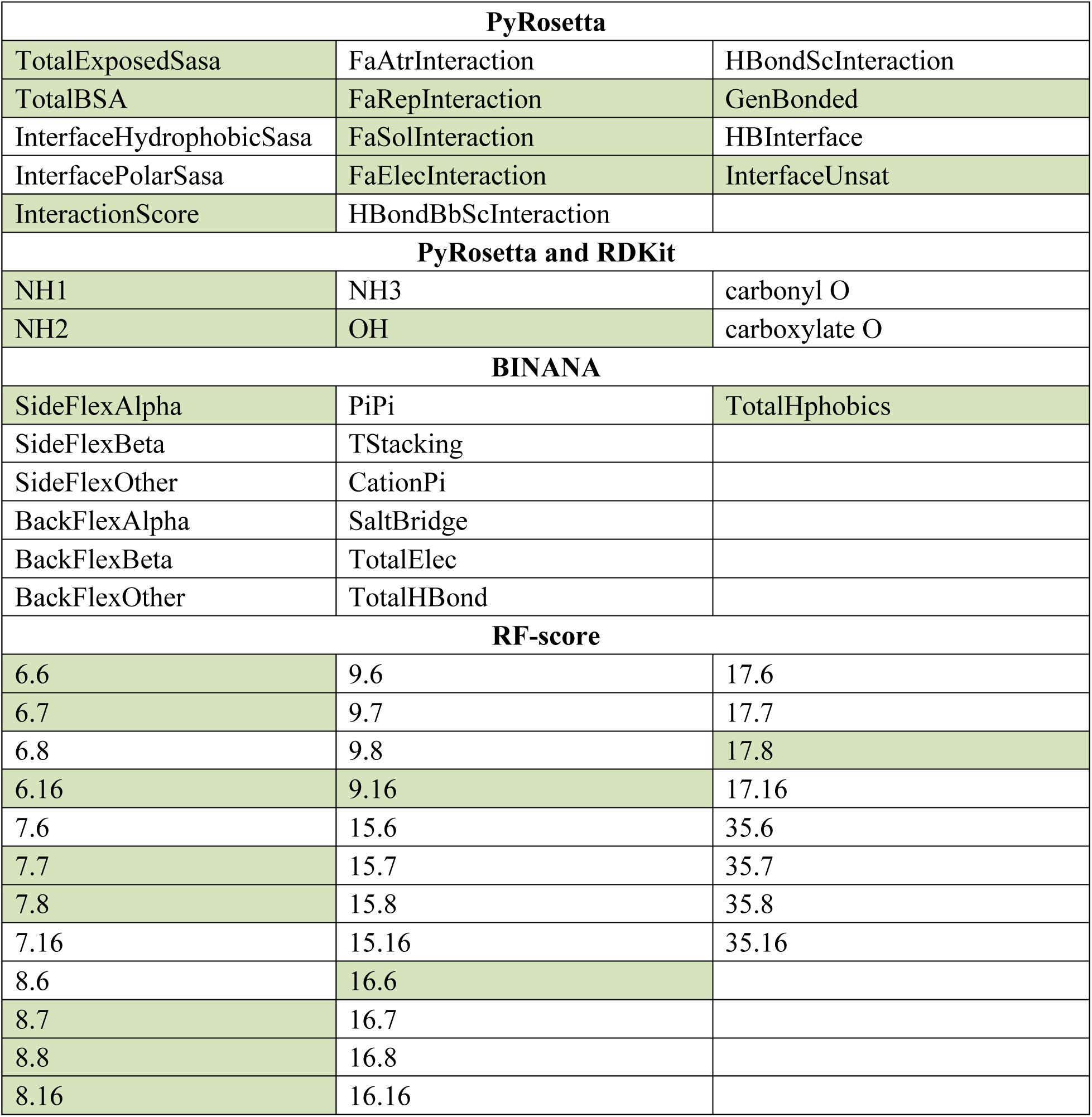
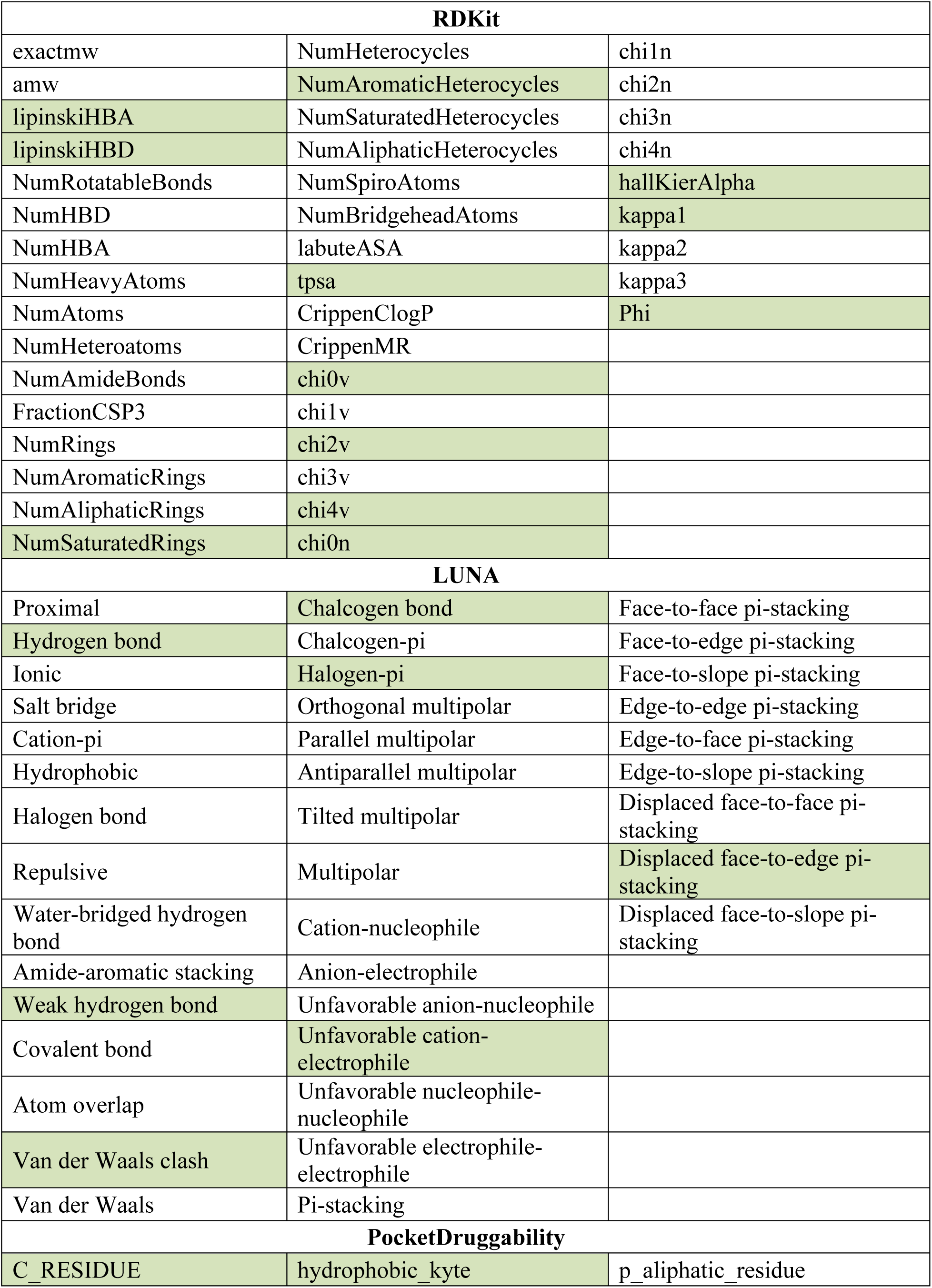
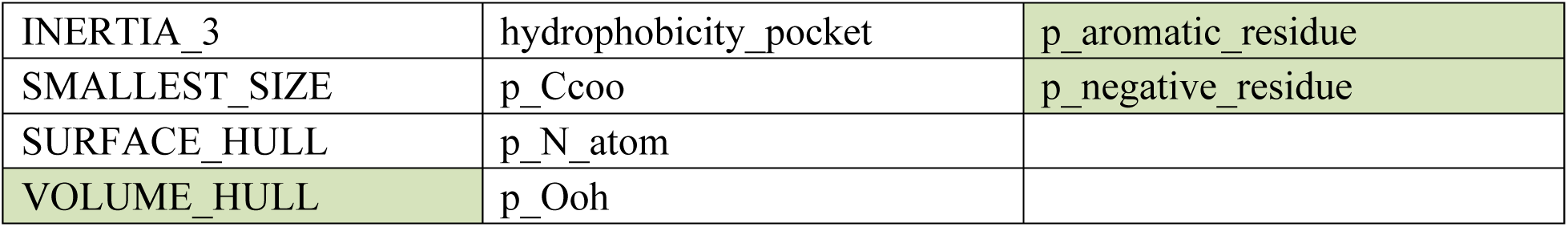
All features calculated for new version of vScreenML. Green indicates the most important features for inference.

**Table S3.**
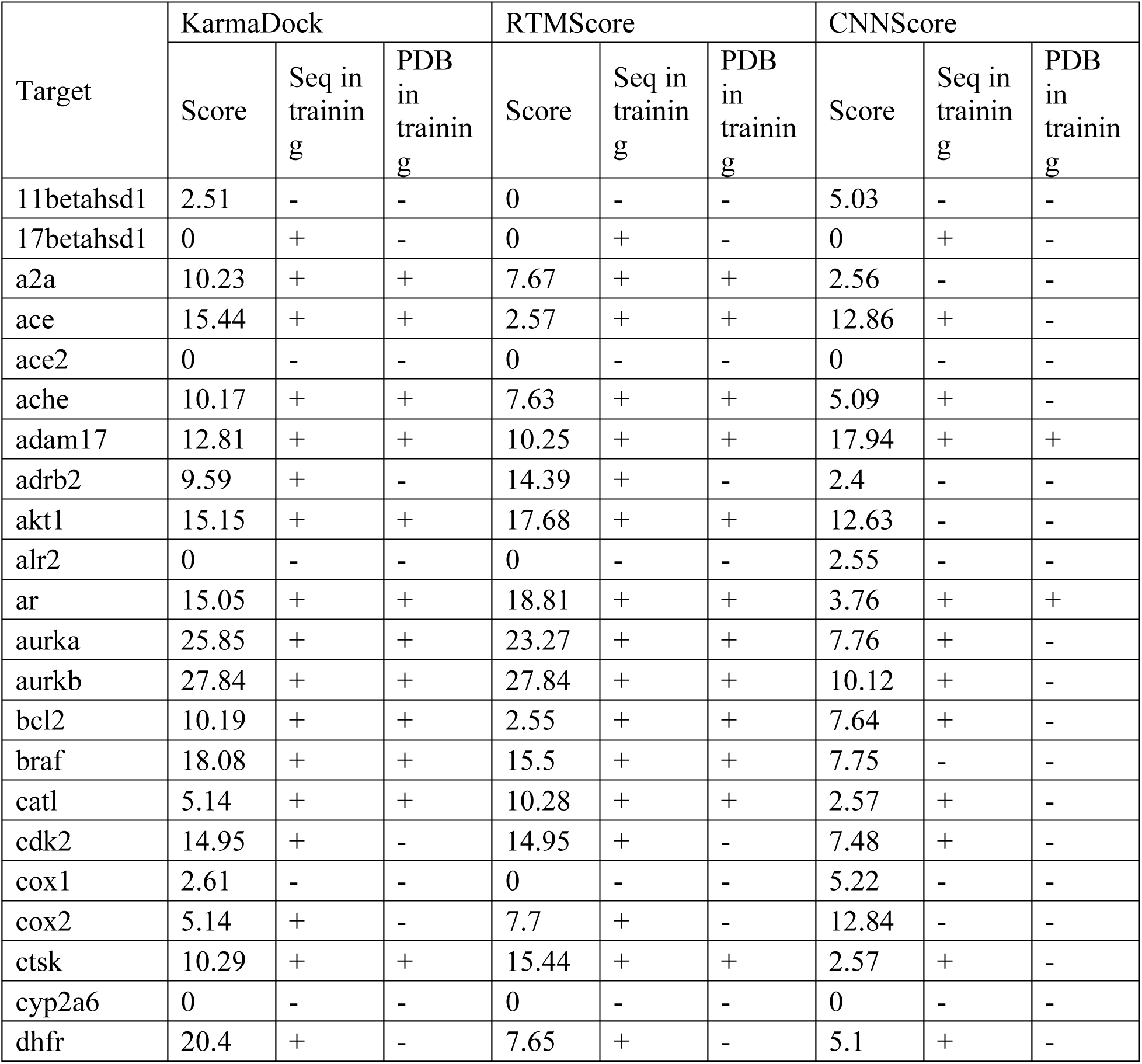
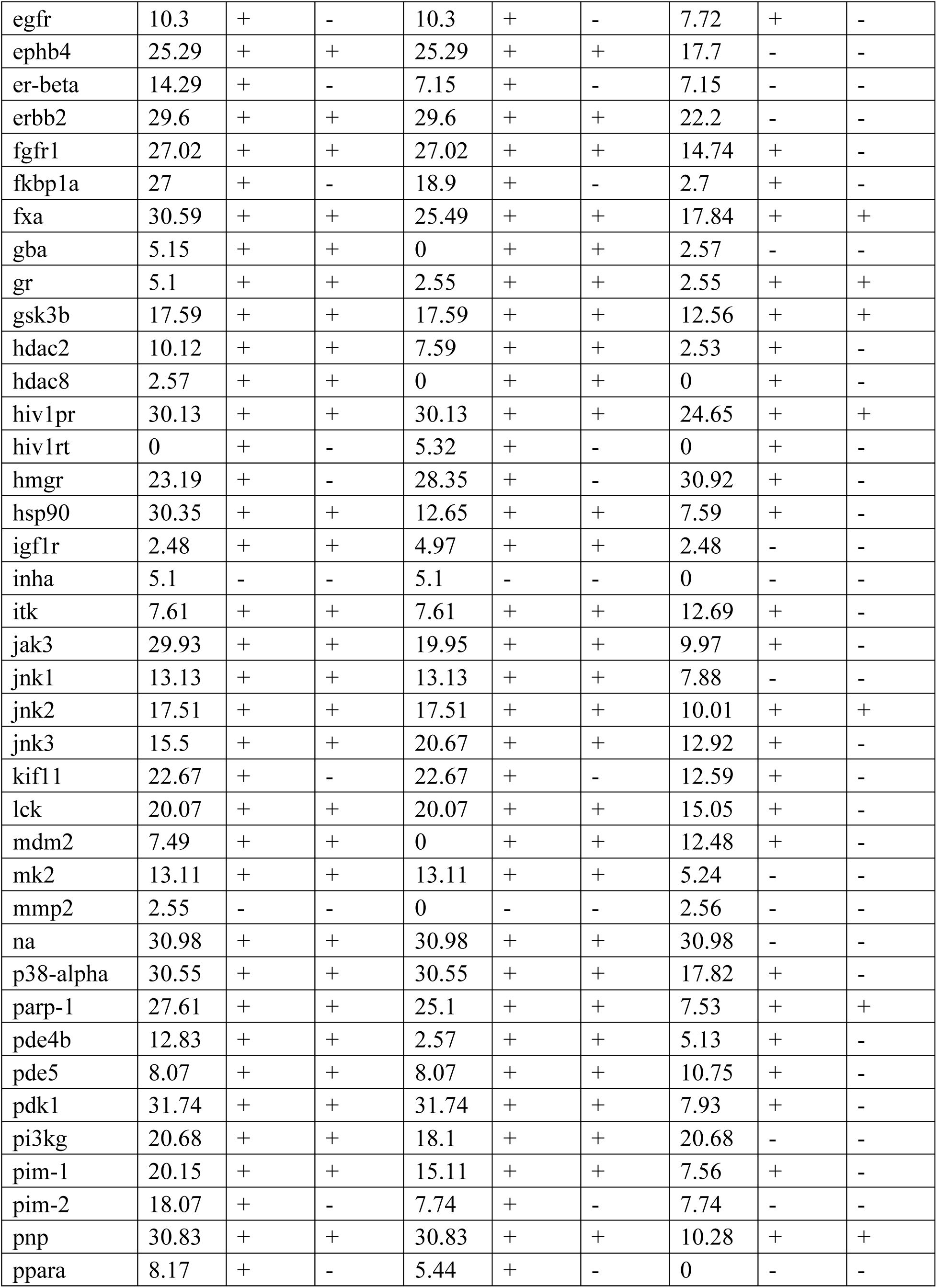
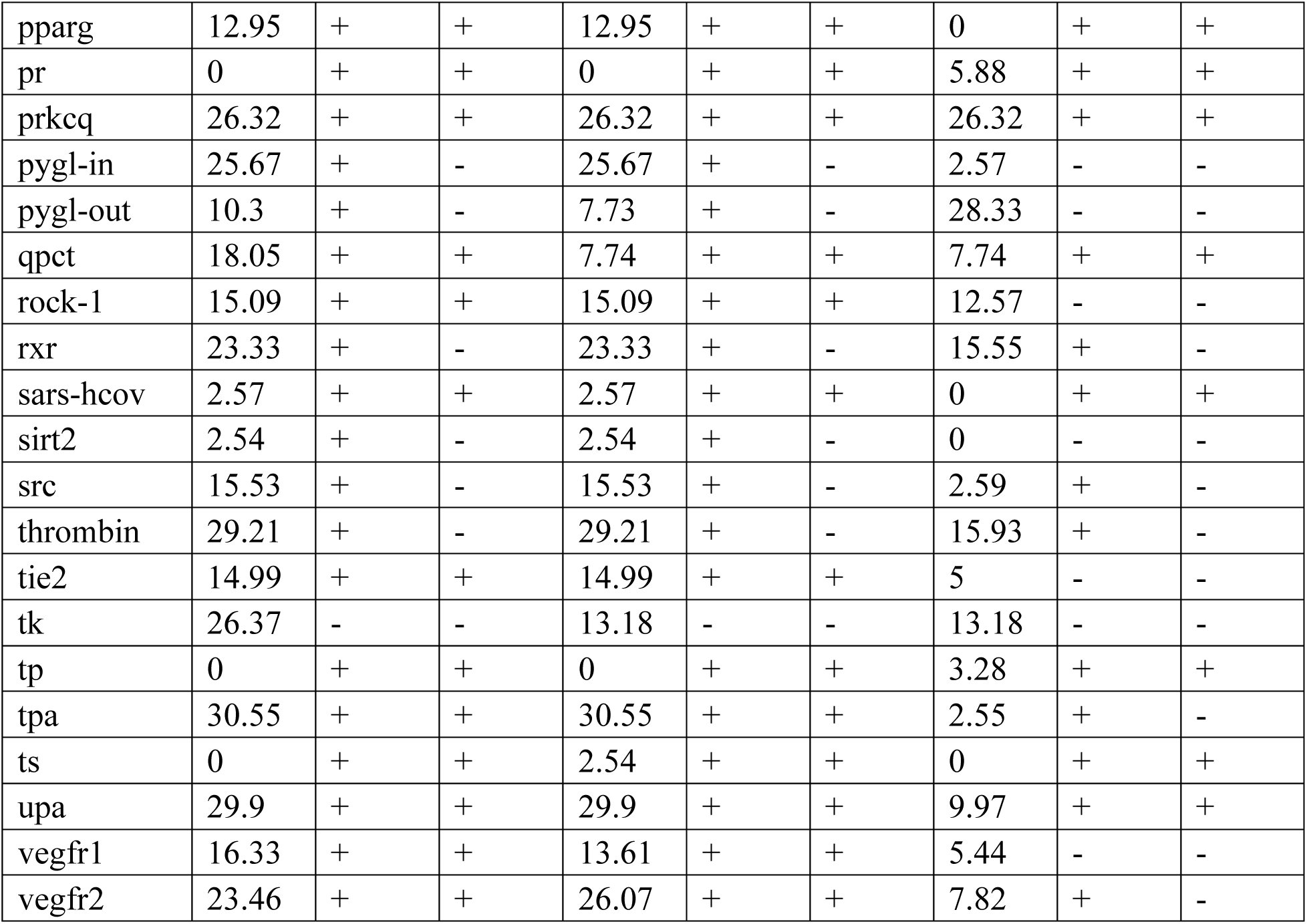
EF1% ranking performance of KarmaDock, RTMScore and CNNscore of gnina on pre-docked DEKOIS2 data set.

